# Small molecule oxybutynin rescues proliferative capacity of complex III-defective MPCs

**DOI:** 10.1101/2024.11.08.622694

**Authors:** Yue Qu, Kaydine Edwards, Muying Li, Yang Liu, Pei-Yin Tsai, Chloe Cheng, Jamie Blum, Noel Acor, Tenzin Oshoe, Kyra Rooney, Claire Walter, Venkatesh Thirumalaikumar, Anna Thalacker-Mercer, Aleksandra Skyricz, Joeva J Barrow

**Author notes:** Co-first author (Yue Qu, Kaydine Edwards).

## Abstract

Mitochondrial disease encompasses a group of genetically inherited disorders hallmarked by an inability of the respiratory chain to produce sufficient ATP. These disorders present with multisystemic pathologies that predominantly impact highly energetic tissues such as skeletal muscle. There is no cure or effective treatment for mitochondrial disease. We have discovered a small molecule known as oxybutynin that can bypass Complex III mitochondrial dysfunction in primary murine and human skeletal muscle progenitor cells (MPCs). Oxybutynin administration improves MPC proliferative capacity, enhances cellular glycolytic function, and improves myotube formation. Mechanistically, results from our isothermal shift assay indicates that oxybutynin interacts with a suite of proteins involved in mRNA processing which then trigger the upregulation biological pathways to circumvent CIII mitochondrial dysfunction. Taken together, we provide evidence for the small molecule oxybutynin as a potential therapeutic candidate for the future treatment of CIII mitochondrial dysfunction.

## INTRODUCTION

Mitochondrial diseases are a diverse group of genetically inherited disorders that arise due to mutations in mitochondrial or nuclear DNA that result in defective mitochondrial energetic capacity and function (Gormon et. al 2016, Schon et. al 2010, Parikh et. al 2015). Incidence rates of 1 in 5000 individuals have been reported, making mitochondrial disease one of the most commonly inherited genetic disorders (Bénit et. al, 2009) . Most mitochondrial disease mutations culminate in defects that alter the respiratory chain (RC), leading to failure of the oxidative-phosphorylation (OXPHOS) system to generate ATP (Shepherd et. al, 2006). Amongst the RC deficiencies, Coenzyme Q: Cytochrome C oxidoreductase or complex III (CIII) deficiencies are one of the most devastating mitochondrial disorders (Bénitet. al, 2009). Complex III mediates the transfer of electrons from reduced coenzyme Q to cytochrome C and maintains the critical Q cycle (Mulkidjanian, 2010). The complex is composed of 11 protein subunits, three of which are essential for catalytic function: Cytochrome C1, Cytochrome b and the Rieske protein (Ndi et. al, 2018). Clinical cases have identified select genes including *BCS1L* and *MT-CYB* that are known to be associated with CIII deficiency but, to date, there remain genetically unresolved cases (Fernández-Vizarra & Zeviani, 2015, Gill Borlado et. al, 2010) . Complex III mitochondrial dysfunction leads to multisystemic physiological decline due to critical failures to sustain cellular energetic demand (Alston et. al 2017). Specifically, the classical clinical presentation of mitochondrial CIII deficiency include muscle weakness, encephalomyopathy, cardiomyopathy, and ataxia all emerging in early childhood (Mordaunt et. al, 2015). Furthermore, as highly metabolic tissues such as skeletal muscle are usually the first to succumb to mitochondrial dysfunction (Protasoni et. al 2021), once a patient is diagnosed, the prognosis is poor and often results in early death. Currently, there is no effective therapy or cure, and thus, mitochondrial disease warrants extreme investigative focus. Given that mature skeletal muscle tissue is post-mitotic, a promising avenue for therapeutic targeting is to boost skeletal muscle regeneration by improving the metabolic capacity of the muscle progenitor cell (MPCs) pool. If one could boost the regenerative capacity of MPCs in the skeletal muscle of patients with mitochondrial disease, this could allow for the expansion of the stem cell pool and promote enhanced regeneration and repair of the damaged skeletal muscle tissue.

Herein, we report results from a high-throughput small molecule screen in mitochondrial complex III-defective skeletal MPCs the discovery of a small molecule compound known as oxybutynin that can circumvent mitochondrial CIII dysfunction. Oxybutynin administration in both murine and human mitochondrial CIII-impaired MPCs potently rescues the proliferative capacity of these defective cells and enhances cellular glycolytic function. Oxybutynin-mediated improvements at the muscle progenitor cellular stage translates to improved myotube formation with enhanced glycolytic capacity. We further demonstrate using isothermal shift assay that oxybutynin associates with a suite of proteins involved in RNA splicing and mRNA processing which then trigger the upregulation of RNA metabolic processing pathways to improve MPC proliferative capacity. Our results highlight the small molecule oxybutynin as a potential therapeutic candidate for the future treatment of CIII mitochondrial disorders.

## RESULTS

### Small molecule compound oxybutynin rescues proliferative capacity of chemically induced complex III-defective muscle progenitor cells

To identify small molecules that harbor therapeutic potential to rescue bioenergetic defects caused by mitochondrial complex III (CIII) mutations, we chemically induced mitochondrial CIII dysfunction in the murine muscle progenitor cell line (Sol8) via treatment with antimycin (Fig. 1A). Antimycin is a natural product inhibitor that binds to the Qi site of complex III disrupting the Q cycle in the catalytic core of the complex (Zhang et. al, 2021). This disruption prevents the oxidation of ubiquinol in the electron transport chain (ETC) and interferes with the electron transfer from cytochrome b to c (Xiuquan et. al, 2011). This interference impairs respiratory chain (RC) function, thus impacting the ETC’s ability to produce sufficient ATP. As a result, the proliferative capacity of the CIII-defective MPCs is significantly reduced compared to healthy controls, and thus reflects the defective proliferation capacity of MPC’s observed in patients with mitochondrial disease. (Fujimaki et.al, 2017) (Fig. 1B).

**Fig. 1.**
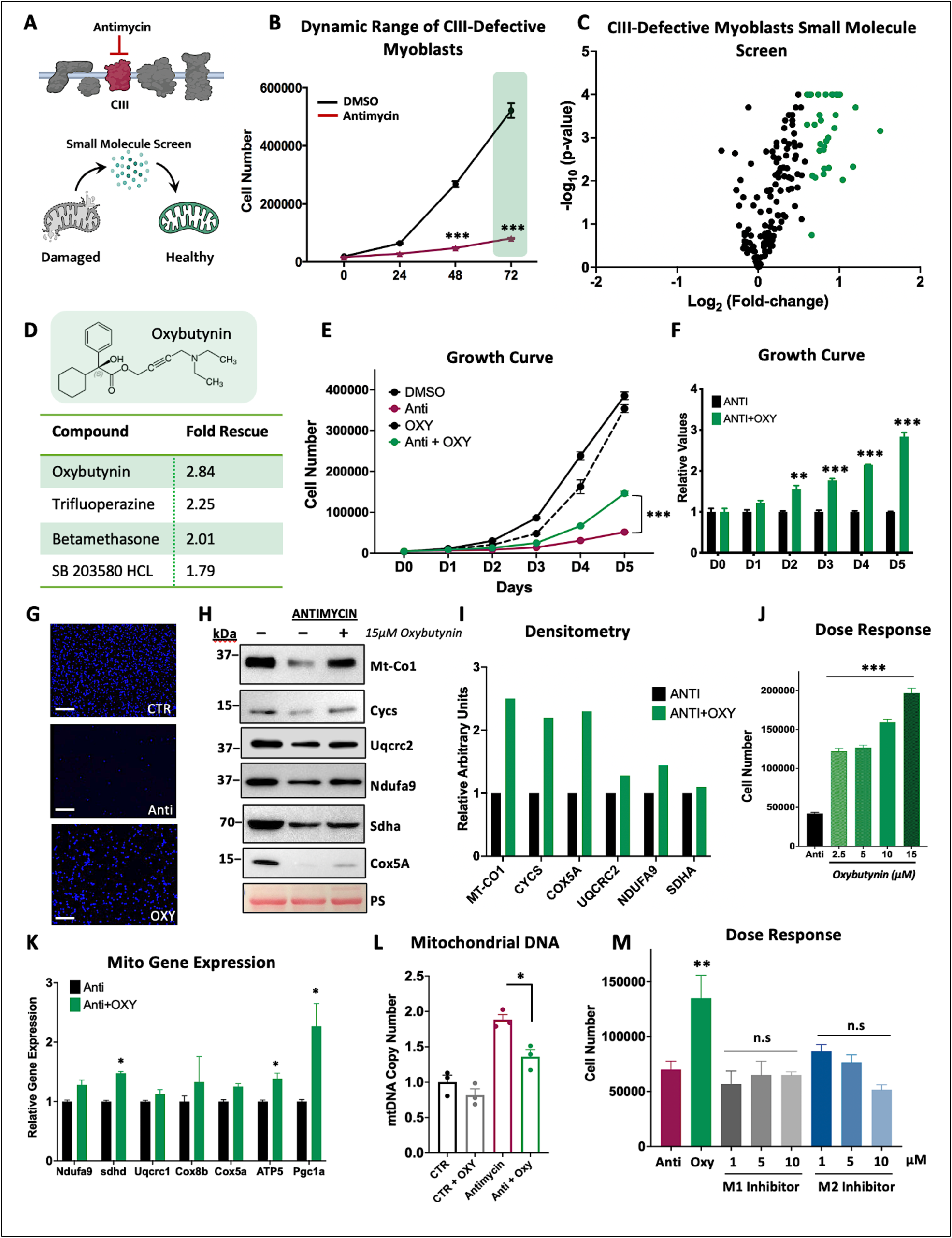
Small molecule compound oxybutynin rescues proliferative capacity of chemically induced complex III-defective muscle progenitor cells. (A). Schematic representation of the chemical inhibition of complex III in Sol8 cells. (B). Proliferation dynamic range between healthy control Sol8 and Sol8 MPCs treated with 100nm of the complex III inhibitor antimycin. (C). Volcano plot of high throughput screen of 50 small molecules. (D). List of top four positive candidates from the screen with oxybutynin exhibiting the most potent rescue potential. (E). Growth curve of healthy Sol8 cells and antimycin-treated Sol8 MPCs treated ± 15μM oxybutynin. (F). Relative quantification of growth curve of complex III inhibited Sol8 MPCs treated ± 15μM oxybutynin. (G). Fluorescent microscopy images of cell number indicated by Hoechst staining (in blue) between healthy Sol8 MPCs and complex III-inhibited Sol8 MPCs ± 15μM oxybutynin. Scale bar=100μm (H). Representative immunoblot of mitochondrial ETC proteins from Sol8 MPCs ± 15μM oxybutynin. Ponceau staining (PS.) is shown as loading control. (I). Densitometry of immunoblot in H. (J). Dose response proliferation rescue of complex III inhibited Sol8 MPCs by oxybutynin. (K). Gene expression of mitochondrial ETC transcripts from antimycin treated Sol8 cells treated ± 15μM oxybutynin. (L). Mitochondrial DNA copy number in complex III inhibited Sol8 cells treated ± 15μM oxybutynin. (M) Growth curve of Sol8 MPCs treated with oxybutynin (15μM) or 10μM of pirenzepine (M1) or methoctramine (M2). All data represent mean ± SEM. ∗ is P-value <0.05, ∗∗ is P-value < 0.01 and ∗∗∗ is P-value < 0.001 by Unpaired Student’s T test.

Leveraging this model system, we performed a targeted small molecule screen of 50 known-bioactive small molecules at varying concentrations to determine if they could rescue the defect in cellular proliferation. The targeted small molecule library was derived from lead hits of a larger parent screen of 10,015 small molecules that had the capacity to increase mitochondrial proteins (Barrow et. al, 2016). Of the 50 small molecules tested, we identified 4 compounds that could increase proliferation of the CIII-defective Sol8 MPCs, with the small molecule oxybutynin exhibiting the most potent rescue potential (Fig. 1C and 1D). Oxybutynin treatment significantly increased CIII-defective MPCs proliferative capacity by 3-fold in a time and dose-dependent manner compared to the DMSO vehicle-treated CIII-defective MPCs (Fig. 1E-G and J). Oxybutynin administration also increased core RC mitochondrial OXPHOS proteins levels compared to untreated controls (Fig. 1H-I). Curiously, this increase was not observed at the transcript level suggesting that the increase in OXPHOS proteins may be due to post-translational regulation (Fig. 1K). We then assessed the levels of mitochondrial DNA in Sol8 MPCs treated with oxybutynin or vehicle controls. As expected, CIII-defective MPCs had higher mitochondrial DNA content as the cell attempts to increase mitochondrial biogenesis to correct the CIII dysfunction (Wallace 2000, Heddi et. al, 1999). Oxybutynin treatment reduced mitochondrial DNA content closer to control levels suggesting improved cellular energetic demands and a reduced need to correct for OXPHOS deficiencies (Fig. 1L).

Oxybutynin is annotated as a non-selective muscarinic acetylcholine receptor inhibitor with higher affinities for the M1 and M2 muscarinic receptor subtypes (McCrery et, al 2006, Noronha-Blob et, al 1991). To determine if the proliferative improvements of the mitochondrial CIII-defective MPCs were occurring via the canonical function of oxybutynin, we treated CIII-defective MPCs with selective antagonists for the M1 (pirenzepine) and M2 (methoctramine) muscarinic receptor subtypes (Lenina et. al, 2022, Alkawadri et. al, 2022) Surprisingly, although oxybutynin treatment continued to improve the proliferative capacity of CIII-defective MPCs, neither the M1 nor M2 muscarinic receptor inhibitors were able to improve cell growth in the face of CIII mitochondrial dysfunction (Fig. 1M). This suggests that oxybutynin is able to bypass CIII mitochondrial dysfunction using alternative pathways. Taken together, these findings identify oxybutynin as a small molecule compound that has the therapeutic potential to bypass CIII-mitochondrial dysfunction and facilitate a significant increase in the proliferative capacity of CIII-defective MPCs.

### Oxybutynin administration improves MPC bioenergetic capacity and proliferation in a genetic mouse model of complex III mitochondrial disease

To ensure that the oxybutynin-mediated rescue of our chemically induced CIII MPCs was not due to a possible chemical interaction with oxybutynin and antimycin, we generated a genetic mouse model of mitochondrial CIII dysfunction. These mice are floxed at exon 2 of the Ubiquinol-cytochrome c reductase, Resike iron-sulfur polypeptide 1 (*Uqcrfs1*) gene and were crossed with a tamoxifen-inducible *Pax7* CRE to create a genetic deletion of *Uqcrfs1* in muscle satellite cells (Fig. 2A). The *Uqcrfs1* gene encodes the Rieske iron-sulfur protein (RISP) which is an essential catalytic subunit of mitochondrial complex III (Saldana-Caboverde et. al, 2020). Ablation of this protein will render complex III inactive, which recapitulates clinical complex III deficiency in human patients (Čunátová et. al, 2024) . As observed in our chemically induced CIII-defective Sol8 MPCs model (Fig. 1B), primary murine MPCs harboring a genetic *Uqcrfs1* deletion in complex III exhibited a significant proliferation defect compared to healthy control *Uqcrfs1^Flox/Flox^* MPCs (Fig. 2B and 1B). Additionally, we observed a marked reduction in the expression of core electron transport chain proteins, similar to the Sol8 MPCs (Fig. 2C and 1H). Furthermore, oxybutynin administration improved the proliferative capacity of the primary murine *Uqcrfs1^-/-^*MPCs compared to vehicle-treated *Uqcrfs1^-/-^* controls, which was consistent with our previous Sol8 MPC observations (Fig. 2D and 2E).

**Fig. 2.**
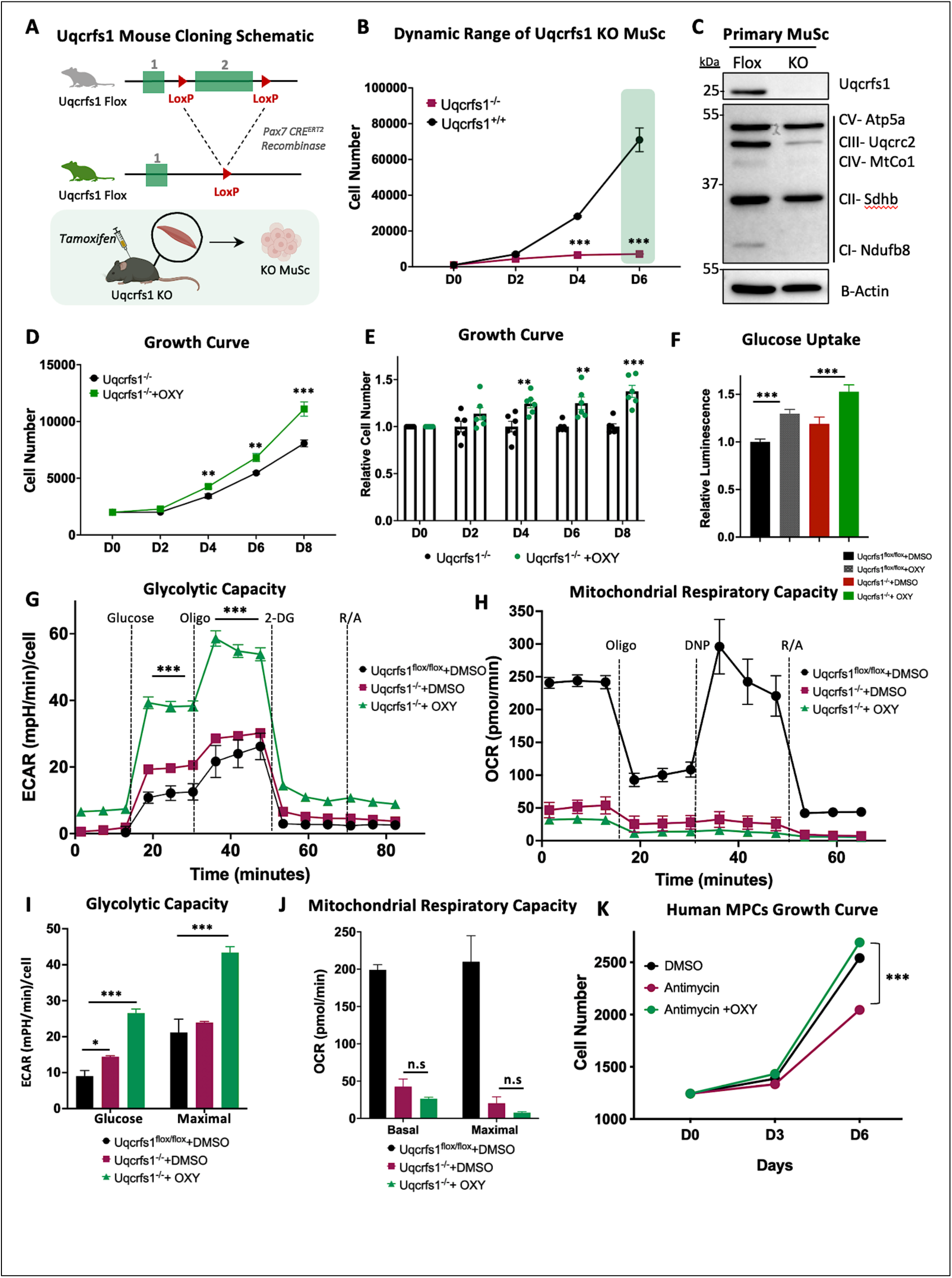
Oxybutynin administration improves MPC bioenergetic capacity and proliferation in a genetic mouse model of complex III mitochondrial disease. (A). Schematic representation of the genetic mouse model of complex III deficiency. (B). Growth curve showing the dynamic range between healthy Uqcrfs1^Flox/Flox^ MPCs with intact complex III and complex III defective Uqcrfs1^-/-^ MPCs. (C). Representative immunoblot illustrating ETC protein expression in the control Uqcrfs1^Flox/Flox^ vs Uqcrfs1^-/-^ knockout MPCs. (D,E). Growth curve of Uqcrfs1^-/-^ MPCs treated ± 10μM oxybutynin. (F). Glucose uptake of Uqcrfs1^Flox/Flox^ and Uqcrfs1^-/-^ MPCs treated ± 10μM oxybutynin. Extracellular acidification (G) and oxygen consumption rates (H) of Uqcrfs1^Flox/Flox^ and Uqcrfs1^-/-^ MPCs treated ± 10μM oxybutynin. (I). Quantification of G. (J). Quantification of H. (K) Growth curve of antimycin (100nm) complex III-inhibited human MPCs treated ± 10μM oxybutynin. All data unless otherwise noted represent mean ± SEM. ∗ is P-value <0.05, ∗∗ is P-value < 0.01 and ∗∗∗ is P-value < 0.001 by Unpaired Student’s T test or two-tailed ANOVA.

To determine if oxybutynin treatment increased cellular growth through improvements in cellular bioenergetic capacity of the primary *Uqcrfs1^-/-^*MPCs, we examined mitochondrial respiratory and glycolytic capacities in *Uqcrfs1^-/-^*MPCs treated with and without oxybutynin. Oxybutynin treatment significantly increased glucose uptake in *Uqcrfs1^-/-^* MPCs compared with untreated *Uqcrfs1^-/-^* controls (Fig. 2F). This increase in substrate availability was associated with an increase in extracellular acidification rate (ECAR) which is a proxy for glycolytic capacity at both basal and maximal glycolytic rates (Fig. 2G and 2I). Oxybutynin treatment did not improve oxygen consumption rates at basal or maximal mitochondrial respiratory capacities in *Uqcrfs1^-/-^* MPCs (Fig. 2H-J) indicating that the mitochondria remained defective. To determine if the oxybutynin-mediated rescue of cellular proliferation in murine MPCs could translate to human cells, we treated mitochondrial CIII-defective human primary MPCs with oxybutynin or vehicle controls and observed a significant improvement in proliferative capacity, indicating that the rescue potential of oxybutynin may be conserved between mice and humans (Fig. 2K). Collectively, these findings suggest that oxybutynin administration rescues mitochondrial complex III-defective MPC proliferation capacity by restoring ATP production through improved glucose metabolism.

### Oxybutynin treatment enhances differentiation capacity of CIII-defective primary MPCs

To determine if the oxybutynin-mediated increase in proliferative capacity of CIII-defective primary MPCs would lead to improvements in myotube formation, we differentiated primary *Uqcrfs1^Flox/Flox^ and Uqcrfs1^-/-^* MPCs in the presence or absence of oxybutynin. Oxybutynin treatment significantly increased myosin heavy chain protein levels and enhanced myotube formation in *Uqcrfs1^-/-^* treated MPCs compared to *Uqcrfs1^-/-^* untreated controls (Fig. 3A and B). Oxybutynin treatment also promoted trending increases in core OXPHOS proteins with no significant changes at the transcript level similar to what was observed in MPCs, suggesting that oxybutynin may operate by regulating post-transcriptional or post-translational processes (Fig. 3C and D). To determine if the enhanced differentiation capacity led to corresponding improvements in cellular bioenergetic capacity, we assessed mitochondrial and glycolytic flux using the Seahorse Bioanalyzer. There were no improvements in mitochondrial respiratory capacity similar to what was observed at the MPC level (Fig. 3E and Figure 2H). Significant increases in cellular glycolytic capacity however were observed which is consistent with the increases seen in MPCs (Fig. 3F-G and Fig. 2G). Taken together, these data suggest that oxybutynin treatment of *Uqcrfs1^-/-^* MPCs harboring CIII mitochondrial dysfunction increases ATP production via improvements in glycolytic bioenergetic capacity to support the energy requirement for enhanced cell growth. The increase in energy production, in turn, promotes improved differentiation into myotubes with sustained elevations in glycolytic capacity compared to *Uqcrfs1^-/-^*untreated controls.

**Fig. 3.**
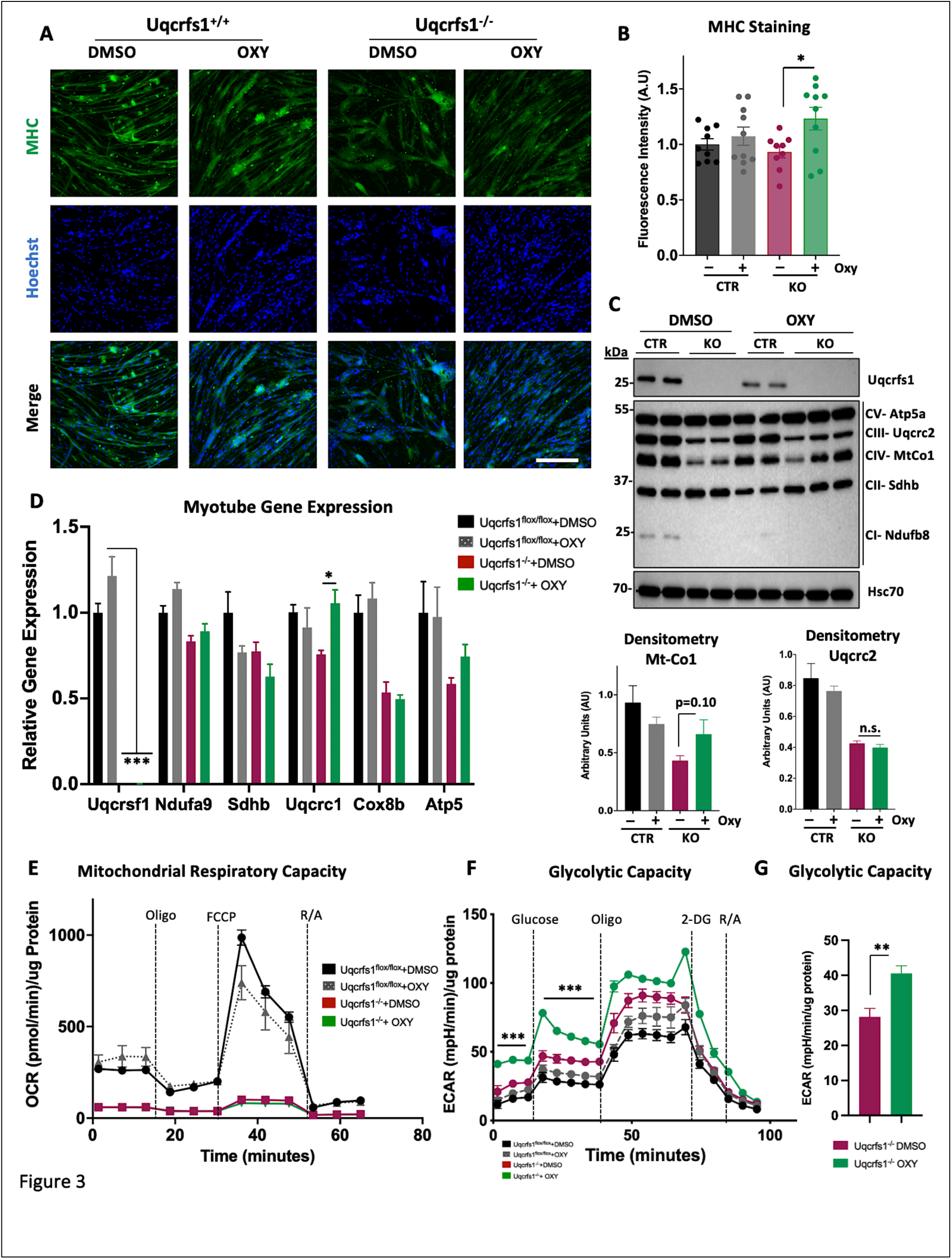
Oxybutynin treatment enhances differentiation capacity of CIII-defective primary MPCs. (A) Fluorescence images of control Uqcrfs1^Flox/Flox^ and knockout Uqcrfs1^-/-^ differentiated myotubes ± 10μM oxybutynin. MHC-Myosin heavy chain. (B). Quantification of the MHC fluorescence of myotubes in A. (C). Representative immunoblot illustrating ETC protein expression in the control Uqcrfs1^Flox/Flox^ and knockout Uqcrfs1^-/-^ differentiated myotubes ± 7.5μM oxybutynin (top). Densitometry of mtCo1 and Uqcrc2 protein expression (below). (D). Relative gene expression of ETC genes in the control Uqcrfs1^Flox/Flox^ and knockout Uqcrfs1^-/-^ differentiated myotubes ± 7.5μM oxybutynin. Oxygen consumption (E) and extracellular acidification rates (F) of control Uqcrfs1^Flox/Flox^ and knockout Uqcrfs1^-/-^ differentiated myotubes ± 7.5μM oxybutynin. (G) Quantification of glycolytic capacity from F. All data unless otherwise noted represent mean ± SEM. ∗ is P-value <0.05, ∗∗ is P-value < 0.01 and ∗∗∗ is P-value < 0.001 by Unpaired Student’s T test or two-tailed ANOVA.

### Oxybutynin enhances the RNA transcription machinery in Uqcrfs1^-/-^ MPCs

As previously shown, oxybutynin treatment bypasses CIII mitochondrial dysfunction in Sol8 MPCs independent of its canonical role as a muscarinic receptor antagonist (Fig. 1M), however, the mechanism of how this is achieved is unknown. To gain mechanistic insight, we first wanted to identify the target protein binding partners for oxybutynin. We therefore performed an isothermal shift assay profiling (iTSA) in Uqcrfs1^-/-^ MPCs treated with or without oxybutynin (Fig. 4A). The iTSA technique captures the differential thermal stability of compound bound proteins compared to unbound proteins and revealed nine proteins that were significantly protected from thermal degradation and thus projected to be bound to oxybutynin (Fig. 4B) (Ball et. al, 2020, Mateus et. al, 2020). Intriguingly, gene ontology revealed that the majority of the binding targets for oxybutynin were all components of the RNA splicing and processing machinery (Fig. 4C). To determine which gene pathways were being upregulated, we analyzed the transcriptome of primary Uqcrfs1^-/-^ MPCs treated with or without oxybutynin. Principal component analysis (PCA) confirmed a distinct treatment response to oxybutynin and transcriptomic analysis revealed several genes that were differentially upregulated in oxybutynin treated Uqcrfs1^-/-^ MPCs compared to vehicle treated Uqcrfs1^-/-^ controls (Fig. 4D and 4E). Notably many genes that were significantly upregulated ≥1.5 fold with oxybutynin treatment belonged to the RNA processing pathway, such as *Snora23, Snora53*, *and Rpl19* which is consistent with the finding that protein binding targets of oxybutynin belong to the RNA-processing protein machinery (Fig. 4F). In addition, several genes belonging to amino acid transport pathways such as the sodium-linked glutamine transporter *Slc1A5* were also upregulated suggesting that oxybutynin may increase the uptake of amino acids towards catabolic pathways as an alternative means to generate ATP similar to what was observed with enhanced glucose uptake leading towards increased rates of glycolysis (Figure 2F and 2G). These findings suggest that oxybutynin’s ability to circumvent CIII mitochondrial dysfunction in MPCs may be a result of the compound’s ability to upregulate the cellular RNA processing machinery and increase amino acid uptake.

**Fig. 4.**
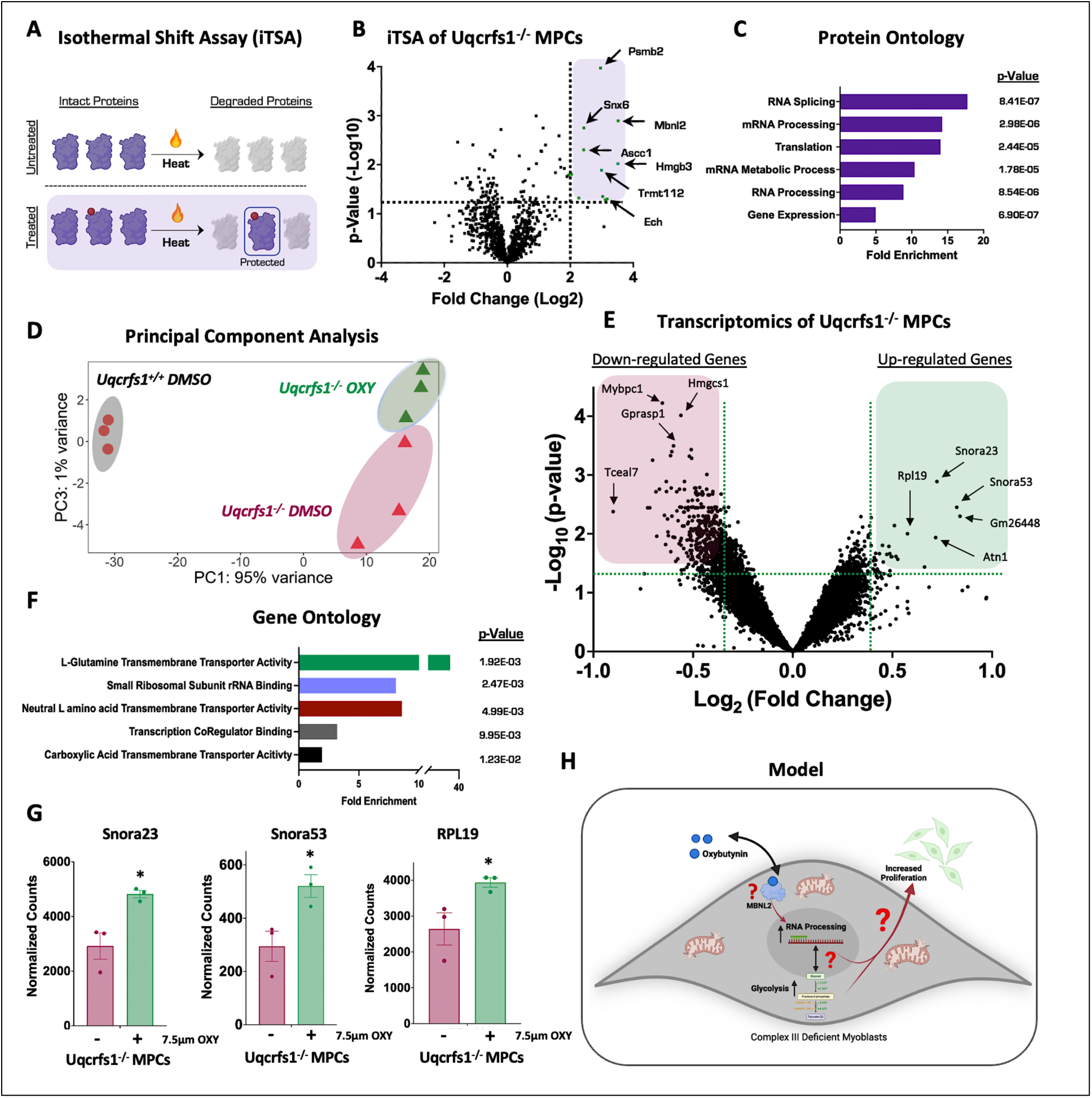
Oxybutynin enhances the RNA transcription machinery in Uqcrfs1^-/-^ MPCs. (A). Schematic representation of the isothermal shift assay (iTSA) (B). Volcano plot of thermal proteome profiling results of Uqcrfs1^-/-^ treated with 10μm oxybutynin compared to vehicle treated Uqcrfs1^-/-^ control MPCs. Putative protein binding partners of Oxy highlighted in purple. (C). Gene ontology from iTSA in B. (D). Principal component analysis of RNA sequencing experimental groups. (E). RNA-Seq Transcriptomic analysis of knockout of Uqcrfs1^-/-^ treated with 7.5μM oxybutynin compared to vehicle treated Uqcrfs1^-/-^ control MPCs. (F) Gene ontology illustrating the top pathways connected to the upregulated genes in E. (G) Representative genes from RNA Seq Analysis. (H). Proposed model for oxybutynin’s mechanism of action in Uqcrfs1^-/-^ MPCs.

## DISCUSSION

Mitochondrial disease is an urgent, incurable health concern with devastating multisystemic pathologies. Of the different mitochondrial disorders, those resulting from mutations affecting the terminal protein complexes, such as Complex III, in the electron transport chain (ETC) are amongst the most severe (Tucker et. al, 2013). Patients with these mitochondrial mutations can no longer keep up with cellular energy demands and experience progressive decline and death of highly metabolic tissues such as the skeletal muscle (Watson et. al, 2020, Mukherjee et. al, 2020).

In this work, we discovered a molecule known as oxybutynin from a small molecule screen that can significantly enhance proliferation in human and murine primary MPCs despite severe CIII mitochondrial dysfunction. This increase in cell growth was both dose and time-dependent and corresponded to an increase in protein expression of other key mitochondrial electron transport chain proteins. In addition, oxybutynin treatment increased the rate of glucose uptake in MPCs to provide additional substrates to fuel the increase in glycolytic activity to produce ATP. Since the improvements in OXPHOS protein expression did not lead to functional rescue via oxygen consumption rate, the upregulation of glycolysis seemed to be more significantly influenced by oxybutynin treatment and thus could be responsible for the improvement in proliferation noted in these mitochondrial complex III defective cells. Finally, oxybutynin administration to MPCs led to an improvement in myotube formation, where the significant upregulation in glycolytic capacity persisted compared to untreated controls.

Oxybutynin belongs to a class of medicinal compounds known as antispasmodics and is used to treat bladder incontinence (Chen et. al, 2024, Lucente et. al, 2011). It functions by competitively binding to muscarinic receptors of the detrusor muscle, where it inhibits acetylcholine stimulation leading to the decrease of muscle spasms of the bladder that would stimulate urination (Jirschele et. al, 2013, Andersson et. al, 2001, Yarker et. al, 1995). However, this molecule remains unexplored in the context of mitochondrial disease. In this work, it does not appear that oxybutynin bypasses CIII mitochondrial dysfunction through its role as an antagonist to the muscarinic receptors. Indeed, when selective inhibitors to the M1 and M2 subclass of muscarinic receptors were administered to Uqcrfs1^-/-^ MPCs, the treatment displayed no rescue in MPC proliferation. It is possible that the oxybutynin-mediated rescue of Uqcrfs1^-/-^ MPCs are derived from off-target effects, however, we believe this is unlikely as the increase in oxybutynin-mediated cell growth were both dose and time dependent, arguing against the possibility that oxybutynin is binding to other therapeutic targets to cause off-target effects. To determine the potential mechanisms of action, we took an unbiased approach to define the binding partners of oxybutynin in Uqcrfs1^-/-^ MPCs. iTSA analysis uncovered nine potential protein binding partners for oxybutynin. Interestingly, muscleblind like protein 2 (MBNL2), a key RNA binding protein involved in modulating alternative splicing of pre-mRNAs (Piasecka et. al 2024, Pascual et. al, 2005) and others involved in the RNA processing machinery were identified. Consistent with these findings, transcriptomic analysis of oxybutynin-treated vs untreated Uqcrfs1^-/-^ MPCs and corresponding gene ontology analysis revealed an upregulation of genes involved in RNA processing. However, how the increase in the RNA processing machinery translates to improvements in complex III defective MPC proliferation requires further investigation. Global transcriptomics analysis also revealed an upregulation of a series of transporters that increase amino acid uptake such as glutamine into the cell which could be used as alternative sources to generate ATP. Taken together, these results point to the possibility that the increased proliferation seen in complex III-deficient MPCs treated with oxybutynin may be a result of oxybutynin increasing the uptake amino acids that can be used as energy substrates.

In conclusion, mitochondrial disease remains an incurable devastating health concern. From a small molecule screen, we uncovered the small molecule oxybutynin that can improve proliferative capacity of mitochondrial CIII-deficient skeletal muscle progenitor cells. Our data suggests that oxybutynin binds, either directly or indirectly, to RNA processing enzymes to increase the expression of the RNA processing machinery as well as the increased expression of a series of amino acid transporters. However, how this upregulation of RNA processing and amino acid transporters leads to the enhanced uptake of circulating glucose to fuel the increase in glycolysis to produce ATP requires further investigation (Fig. 4G).

## MATERIALS AND METHODS

### Animal Models

All animal experiments were approved by the Institutional Animal Care and use Committee at Cornell University. C57BL/6J *Uqcrfs1^fl/fl^*(*RISP^fl/fl^*) mice were a gift from Dr. Paul Schumacker at Northwestern University and have previously been described (Sena et. al, 2013; Weinberg et. al, 2019). B6.Cg-Pax7 tm1(cre/ERT2)Gaka /J (#017763) mice were obtained from Jackson Laboratory. Briefly, to achieve murine MPC specific deletion of *Uqcrfs1*, mice homozygous for floxed *Uqcrfs1* were crossed with mice heterozygous for *Pax7*^Cre/ERT2^ to generate progeny expressing both homozygous floxed *Uqcrfs1* and heterozygous *Pax7*^Cre/ERT2^. At 3 weeks, these mice were intraperitoneally injected with tamoxifen (Sigma Aldrich, T5648-1G) or sunflower oil vehicle (for the control group) for 5 days at a concentration of 30mg/g of bodyweight, followed by a 7-day washout period. After the washout, mice were euthanized using CO_2_ asphyxiation and muscle tissue was harvested from both hind limbs from each mouse as previously described (Qu et. al, 2023) . All mice were maintained under 12hr light and dark cycles and fed ad libitum with standard irradiated rodent chow.

### 2.1 High-throughput chemical screening

105 lead compounds from a previous mitochondrial complex I deficient cell-based screen of 10,015 chemical compounds were cross screened in chemically induced mitochondrial complex III deficient muscle progenitor cells (Barrow et. al, 2016) . Briefly, Sol8 MPCs (ATCC CRL-2174) were seeded at 7.5×10^3^ per well in collagen-coated 12-well plates in high glucose (4.5 g/L) DMEM medium supplemented with 20% FBS, 1% Pen/Strep, 25mM HEPES (HG Medium) and incubated for 24hr at 37°C and 5% CO_2_. Cells were then washed with PBS and switched into low glucose media (DMEM with 1g/L D-glucose, supplied with 20% FBS, 1% P/S, 25mM HEPES, 1mM Pyruvate, 200uM Uridine) and subjected to treatment with varying concentrations of lead small molecule compound, including oxybutynin (AstraZeneca-AB7701) or vehicle control with or without antimycin (Sigma-A8674) for 72hrs. At the end point, cells were briefly rinsed in PBS and stained with Hoechst at a concentration ratio of 1:2000 (Invitrogen H21492) and propidium iodide at a concentration ratio of 1:500 (Invitrogen P3566) in FBS-free low glucose medium for 30 mins. MPC concentration was then assessed using the Celigo S Image Cytometer automated

microscopy platform (Nexcelom).

### 2.2 Primary Human and Mouse MPCs Isolation

Primary human MPCs cells were obtained from Dr. Anna Thalacker-Mercer at the University of Alabama and cultured following a previous published protocol (Gheller et. al, 2021; Qu et. al, 2023). Briefly, cells were seeded at 3×10^3^ per well in collagen-coated 96-well plates in human cell growth medium (hGM; 79% Ham’s F12, 20% fetal bovine serum, 1% penicillin/streptomycin, 5 ng/mL basic fibroblast growth factor [bFGF], pH 7.4, passed through a 0.22 μm filter) for 24hr at 37°C and 5% CO_2_. Cells were then treated with hGM with 100nM antimycin and 10uM oxybutynin with the appropriate controls as indicated in the Fig.s. The hGM along with antimycin and oxybutynin was refreshed every 48hr to maintain cell growth. On day 9, cells were stained with Hoechst and PI and imaged for cell proliferation as described above. Primary murine Uqcrfs1 Flox and KO MPCs were obtained from 4-week-old male CB57B/6J *Uqcrfs1^fl/fl^* mice treated with or without Tamoxifen according to published protocols (Qu et. al, 2023) . Briefly, muscle tissue of both hindlimbs were mechanically sheared and washed with HBSS (Invitrogen, 14170-112). The muscle tissue was then resuspended in 4mL of digest media (1.2Uml Dispase II (Roche 295-825) and 1800 U/mL collagenase II (Gibco, 17101-015) in 2.5mM CaCl_2_ in HBSS) and incubated at 37°C shaking water bath for 45mins. Following incubation, 4mL of D10 medium (DMEM, 10% FBS and 1% P/S) was added and then passed through a 100uM sterile filter. The flow-through was then filtered through a 40uM sterile filter. Cells were then centrifuged at 800x g for 5 mins. Cells were washed with 4mL of D10 and centrifuged to pellet the cells. The cell pellet was then resuspended in 5mL of primary MPCs growth medium (PMGM) (20% Horse serum, 1% Pen/Strep, and 25ng/ml bFGF) and plated in a T-25 flask coated in 2% gelatin incubated at 37°C with 5% CO_2_. After 48 hrs of incubation, the PMGM were collected and purified using negative selection.

### 2.3 Genetic Knockout using Adenovirus CRE

A total of 4x10^5^ primary *Uqcrfs1^fl/fl^* MPCs were seeded in 2% gelatin coated T75 flask and incubated at 37°C with 5% CO_2_ for 24 hrs. Following seeding, cells were treated with commercial adenovirus GFP (Ad4627, U of Iowa-VVC) or CRE (Ad4697 U of Iowa-VVC) at an MOI of 250 per flask. Treated flasks were incubated for 72hrs to allow for complete infection by the virus. Genetic knockout of *Uqcrfs1* was confirmed via western blot and downstream experiments were conducted as indicated.

### 2.4 Primary Murine MPCs Treatments

MPCs culture and differentiation was performed as previously described (Qu et. al, 2023, Earle et. al, 2020). Briefly, *Uqcrfs1* GFP treated control and CRE treated knockout primary MPCs were seeded at a density of 9.5x10^4^ cells per well in 2% gelatin coated 24-well plates, and 1.6x10^4^ cells per well in 2% gelatin coated seahorse plates. Cells were seeded in F10 primary MPCs growth media supplemented with 20% horse serum, 1% penicillin/streptomycin and bfgf (R&D Systems, 233-FB) at a concentration ratio of 25ng/mL. Adenovirus GFP and CRE treated MPCs were seeded in 2% gelatin coated T75 flasks at a concentration of 4.0x10^5^ cells/flask. 24 hours after seeding cells were treated with 7.5μM of oxybutynin (AstaTech, AB7701), 1-10μM pirenzepine (Sigma), 1-10μM methoctramine (Sigma), or DMSO vehicle and incubated for 5 days at 37°C. For differentiated myotubes, cells were seeded in 24-well plates as described above. 24 hours after seeding, F10 media was replaced with IMDM (Gibco, Ref12440-053) media supplemented with 2% horse serum. Cells were then treated with 7.5μM of oxybutynin or with DMSO vehicle control. Differentiation media was refreshed every day for 6 days with oxybutynin being refreshed every two days. Agrin (R&D Systems, 550AG/CF) was added at a concentration ratio of 0.1mg/mL to the differentiation media on the third day following seeding up until the 6^th^ day of differentiation. Differentiation was terminated on day 6 and downstream analysis by western blot, qPCR and seahorse were conducted as indicated.

### 2.4 Immunoblotting

Western blot was performed as previously described (Liu et. al, 2023) . Briefly, cells were harvested in RIPA buffer (10 mM Tris-HCl pH 8.0, 1 mM EDTA, 1% Triton X-100, 0.1% Sodium Deoxycholate, 0.1% SDS, 140 mM NaCl, 1X protease and phosphatase inhibitor cocktail) and proteins were quantified using the BCA assay (Pierce 23228). 20 µg of protein was resolved on a 12% SDS-PAGE gel. The following antibodies were used for western blot analysis: Total OXPHOS Rodent WB Antibody Cocktail 1:1000, Uqcrfs1 1:1000, MtCo1 1:2000, Cycs 1:1000, Uqcrc2 1:1000, Ndufa9 1:1000, Sdha 1:10,000, Cox5a 1:1000, and B actin 1:1000 . Corresponding secondary antibodies: Goat Anti-Mouse IgG (H + L)-HRP Conjugate and Goat Anti-Rabbit IgG (H + L)-HRP Conjugate 1:3000. Immunoreactivity was visualized using enhanced chemiluminescence (ECL) substrate (Thermo, 1863096 and 1863097). Immunoblots were imaged using FluorChem imaging system. Images were quantified with densitometry using Fuji (ImageJ2 v2.9.0).

### 2.4 Gene Expression

Gene expression for qRT-PCR and PCR analysis were performed as previously described (Liu et. al, 2023). Total RNA was isolated with Trizol (Invitrogen, 15596-026). Two micrograms of RNA were used to generate complementary DNA (cDNA) with a High-Capacity RNA-to-cDNA™ Kit (Applied Biosystems, 4368813) following the manufacturer’s protocol. For gene expression analysis, cDNA samples were mixed with PerfeCTa® SYBR® Green FastMix® Reaction Mixes (QuantaBio, 95072) and were analyzed by a CFX 384 Real-Time system (Bio-Rad). All primers and sequences are listed in Table 1-1 and available upon request.

**Table 1.1.**
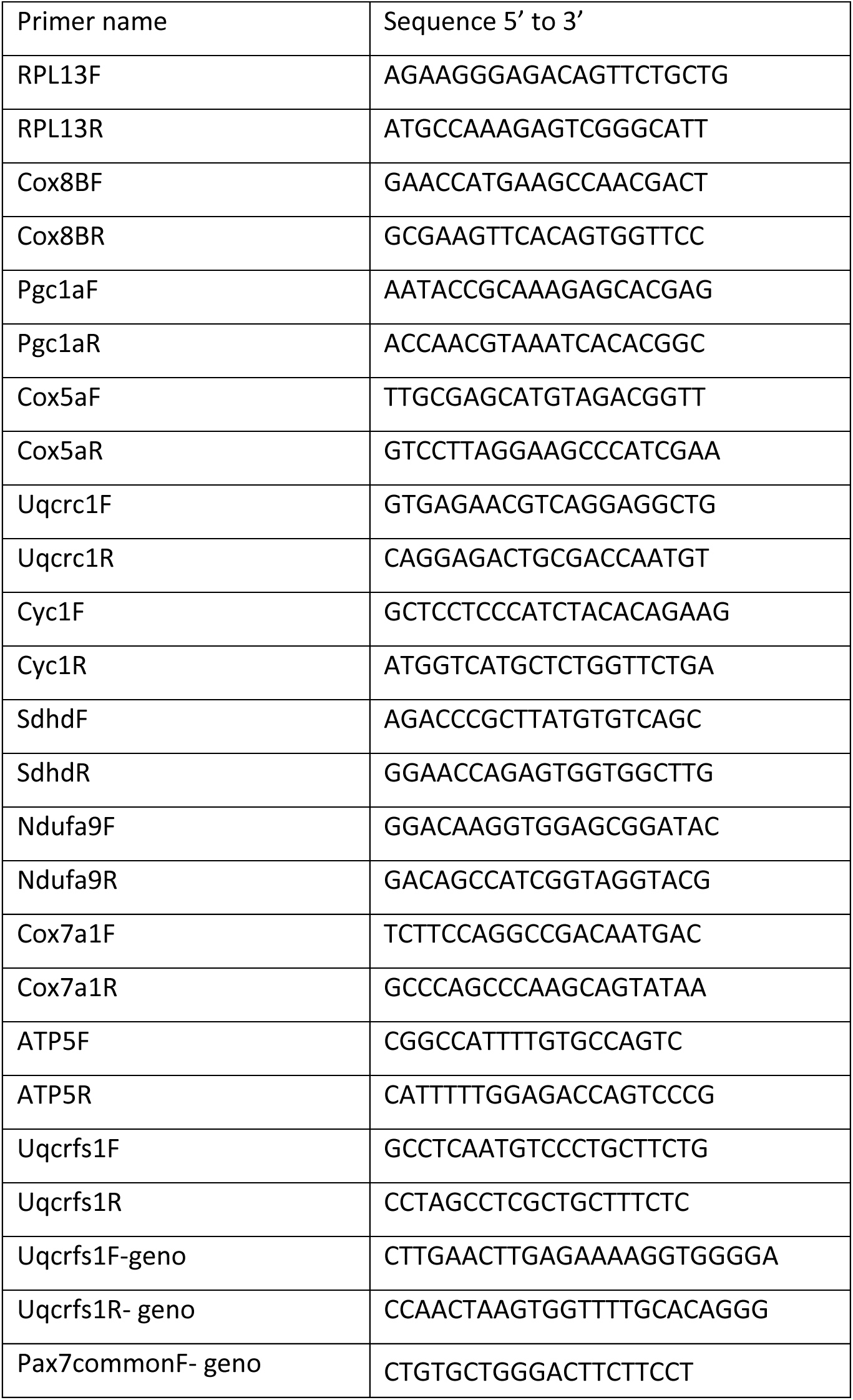
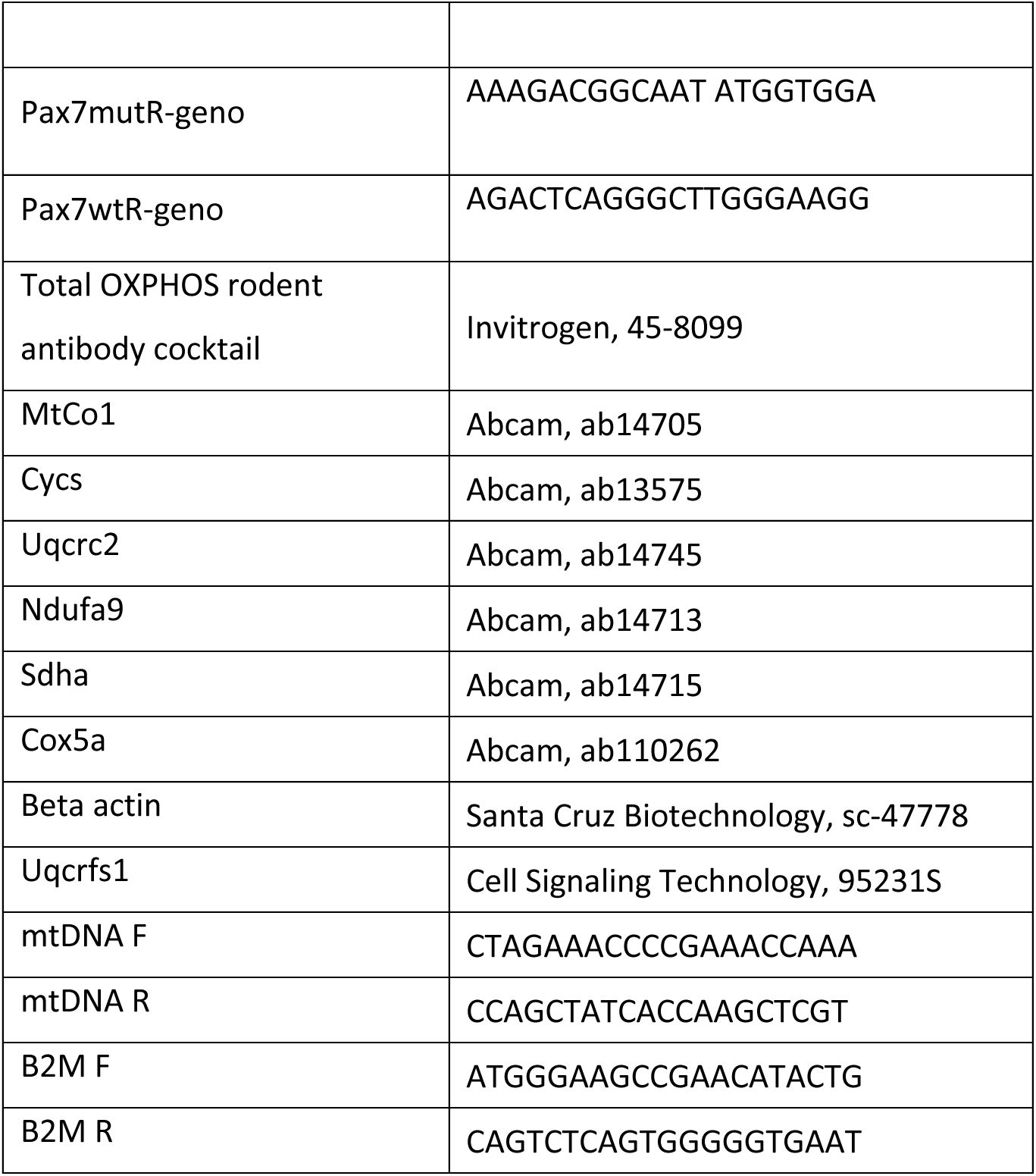
Primer sequences and antibodies catalogue numbers.

### 2.5 Cellular Respirometry

Cellular respirometry was conducted as published previously (Liu et. al, 2023). Treated primary murine *Uqcrfs1* Flox and KO MPCs and myotubes (6.5×10^4^ cells/well) along with the appropriate vehicle controls were seeded in 2% gelatin-coated XF-24 Seahorse plate (Seahorse Biosciences, 102340) and incubated at 37°C with 5% CO_2_ for 24hrs. Following incubation, growth medium was removed, and cells were washed with pre-warmed unbuffered DMEM without bicarbonate (Sigma, D5030) supplemented with 4 mM glutamine. For the mitochondrial stress assay, 1mM sodium pyruvate was added in and pH was adjusted to 7.4 prior to the start of the assay. After the wash, 500 uL of the same buffer was added and cultured at 37°C in a non-CO_2_ incubator for 15 mins. For the mitochondrial stress assay, the Seahorse analyzer cartridge was prepared as follows (final concentration): 1 μM oligomycin, 1.5mM dinitrophenol (DNP), and a mix of 3.75 μM antimycin and 3.75 μM rotenone to each cartridge port. For the glycolysis stress assay, the Seahorse cartridge was prepared as follows (final concentration): 10mM Glucose, 1μM oligomycin, 50mM 2-DG, and a mix of 3.75 μM antimycin and 3.75 μM rotenone to each cartridge port. Oxygen consumption rates (pmole/min) and extracellular acidification rate (mpH/min) were then measured for each treatment condition at 37°C using the Seahorse Bioanalyzer instrument. After measurement, cells were stained with Hoechst and measured by Celigo S Image Cytometer (Nexcelom) to normalize values.

### 2.6 Glucose uptake assay

Primary *Uqcrfs1* Flox and KO MPCs were seeded in 2% gelatin coated 6-well plates and treated with 7.5μM oxybutynin and appropriate vehicle controls for 72 hours according to the indicated experimental design, then 1x10^4^ cells from each group were seeded in a 2% gelatin-coated 96-well plate and incubated at 37°C with 5% CO_2_ for 24 hours. Following incubation, the glucose uptake was measured following the instruction of Glucose Uptake Glo assay kit (Promega, 76201-556).

### 2.7 Isothermal Shift Assay

The isothermal shift assay protocol was modified from (Ball et. al, 2020) . Primary *Uqcrfs1* Flox and KO MPCs (2.0x10^5^ cells) were cultured in 2% gelatin-coated T25 flasks in low glucose (1g/L) DMEM F10 media supplemented with 25ng/mL bFGF until 80% confluency. Cells were then lysed with digitonin lysis buffer [5% digitonin, 50mM Tris pH 7.8, 150mM NaCl supplemented with protease and phosphatase inhibitors] to extract proteins. Protein concentration was measured using the standard Bradford method for protein quantification (Biorad, Hercules, CA, US). MPCs cell lysates were then exposed to 7.5μM oxybutynin or vehicle control at room temperature (RT) for 30 min. Following incubation, cell lysates were diluted with the modified lysis buffer [50mM Tris pH 7.8, 150mM NaCl] to 1 mg/ml. Volume of 100 µl corresponding to roughly 100 µg of protein was aliquoted equally into microcentrifuge tubes and heated at 53°C for 3 min, which corresponds to the reported average meltome of a mammalian proteome (Jarzab et. al, 2020). Samples were then centrifuged for 10 min at 20,000 x g at 4°C and the supernatant (soluble proteins) was transferred to a new tube. Proteins were subsequently precipitated with 2.5x volume of precooled acetone, O/N in the -20°C. Protein pellets obtained after 10 min centrifugation at 20,000 x g were washed with 500 µl of 100 % Methanol (Mass Spec – grade), followed by trypsin digestion and desalting protocol described in (Thirumalaikumar et. al, 2023).

### In-solution trypsin digestion and peptide desalting

Acetone precipitated protein pellets were subjected to protein extraction protocol as described in (Thirumalaikumar et. al,. 2023). Briefly, protein pellets were dissolved in a denaturation buffer (6 M urea, 2 M thiourea dissolved in 50mM Ammonium bicarbonate) the protein extracts were reduced by the addition of DTT (100 μM) for 60 min at RT. Alkylation step was done using (300 μM) iodoacetamide for 60 min at dark. Residual iodoacetamide is quenched by additional (100 μM) DTT for 10 min. Finally, proteins were digested using trypsin/Lys-C mixture (Mass Spec Grade, Promega) for 16 h according to the manufacture’s instruction. Digested peptide samples were desalted using C18 sep-pak column plates as described in Thirumalaikumar et. al,. 2023. The eluted peptides were transferred to Eppendorf low bind tubes and then dried using speed vac. The dried peptides were resuspended using resuspension buffer (5% acetonitrile in 0.1% formic acid).

### Liquid chromatography and mass spectrometry

Approximately, 1μg of the peptides were injected for analysis. The peptide mixtures were separated using a nano-liquid-chromatography system (Dionex ultimate 3000) using an acclaim pepmap C18 column. A flow rate of 300 nL/min was used for the complex peptide separation. The solvent A/B gradient were as follows: being isocratic at 3% B for 17 min (including first 10 min for loading and trap column), linearly increasing to 30% B at 125 min, linearly increasing to 40% B at 145 min, keeping at 95% B from 145.1 min to 155 min, shifting back to 3% B in 0.1 min (valve was returned to load position) and holding until 170 min. The peptide samples were sprayed using a nano bore stainless-steel emitter (Fisher Scientific). Peptides were analyzed using an Orbitrap-Exploris-480 MS™ mass spectrometer. Data was collected using a data-dependent acquisition (DDA) mode. Standard mass spectrometer parameters were kept as described in (Henneberg et. al,. 2022). Briefly: positive ion voltage 2.3 kV, ion transfer tube temperature 320 °C; full scan orbitrap resolution 120,000, scan range *m*/*z* 400-1200, RF lens 50%; ddMS2 filters include monoisotopic peak determination for peptide, intensity threshold 2.0e4, charge 2-6, exclude ions ±10 ppm for 120 s after one scan, exclude isotopes; ddMS2 isolation window *m*/*z* 1.4, HCD normalized collision energies 30%, orbitrap resolution 15,000, 100% normalized AGC target, auto maximum injection time. HeLa digests (Pierce, 88329) have been used to monitor the retention time drift and mass accuracy of the LC before and after each experiment.

### Raw data analysis

Raw data were analyzed using the Proteome Discoverer (version 2.5, ThermoFisher scientific) following the manufacturer’s instruction. PD Search was made using an appropriate recent protein database downloaded from uniport. Common contaminants were compiled and added to the search. SEQUEST HT was used to assign the peptides, allowing a maximum of 2 missed tryptic cleavages. In addition, precursor mass tolerance of 10 ppm and a fragment mass tolerance of 0.02 Da, with a minimum peptide length of 6 AAs were selected. Cysteine-carbidomethylation and methionine-oxidation were selected in default modifications. Label-free quantification (LFQ) based on MS1 precursor ion intensity was performed in Proteome Discoverer with a minimum Quan value threshold set to 0.0001 for unique peptides. The 3’ Top N’ peptides were used for area calculation. The normalized protein abundances were used among the samples and values were log2 transformed in Microsoft excel. Protein target candidates were prioritized based on significance and fold induction over oxybutynin and vehicle control and were subjected to gene ontology analysis to reveal linked biological pathways.

### 2.8 Immunofluorescence

Primary *Uqcrfs1* Flox and KO MPCs were seeded in F10 PMGM growth media at a density of 5x10^4^ cells/cm^2^ and cultured for 24 hours at 37°C with 5% CO_2_. Following incubation, growth media was aspirated, and differentiation was initiated as previously described. Differentiated myotubes were fixed with 3.7% PFA for 45 min at RT with gentle shaking. Cells were then washed 3 times with PBST (PBS with 0.3% Triton) at RT for 5 mins each wash. Cells were then permeabilized with 0.3% Triton X-100 for 15 minutes. MPCs were then washed as described above and then blocked with 5% Donkey Serum (DSB) for 60 mins at RT with gentle shaking. Cells were then incubated with primary eMyHC antibody (1:10, F1.652, Cell Signaling) overnight at 4°C, with gentle shaking. The following day, cells were washed 3 times with PBS at RT followed by incubation with the secondary antibody (Thermofisher, A11008) for 2hrs at RT followed by three washes with PBS at RT. Finally, samples are stained with Hoechst (at a concentration of 0.01mg/mL in PBS) for 10 mins at RT before imaging with Agilent Lionheart. Images were processed with Gen5 software (2023).

### 2.9 Mitochondrial DNA Measurements

The genomic DNA was extracted from Sol8 cells using the QIAamp DNA Micro Kit (Qiagen #56304, Germantown, MD, USA) according to the manufacturer’s guidelines. Subsequently, a quantitative PCR was conducted utilizing primer sequences referencing mitochondrial DNA (mtDNA) and nuclear DNA (nDNA, B2M), respectively. The mitochondrial DNA copy number was quantified following the comprehensive methodology described by (Rooney et. al, 2015) (1) The difference of cycle threshold: Δ𝐶𝑡=𝐶𝑡 (n𝐷𝑁𝐴) −𝐶𝑡 (mt 𝐷𝑁𝐴). (2) The copies of mitochondrial DNA: 𝑚𝑡𝐷𝑁𝐴=2×2Δ𝐶𝑡.

## DATA AVAILABILITY

All data are available upon request pending standard material and transfer agreements between Cornell and requesting institution.

## ACKNOWLEDGEMENTS

The authors sincerely thank Dr. Paul Schumacker from Northwestern University for providing the Uqcrfs1 Flox mice (Waypa et al, 2024). The authors also thank J.T. from the Sethupathy lab at Cornell University for assistance with the PCA plots. We also sincerely thank the Transcriptional Regulation and Expression facility at Cornell University for conducting RNA sequencing. This work was supported by the Hartwell Foundation awarded to J.B. All schematic images were created with Biorender licensed to Cornell University.

## AUTHOR CONTRIBUTIONS

This project was conceptualized by J.J.B., Y.Q., and K.E; Methodology, J.J.B, Y.Q., K.E., P.T., M.L, Y.L., K.R., J.B., V.T., A. S., A. T.M., T.O., N.A.; Investigation Y.Q., K.E., P.T., M.L., N.A.; Formal analysis J.J.B, Y.Q., K.E; Writing J.J.B, Y.Q., K.E; Visualization J.J.B, Y.Q., K.E., N. A.; Supervision, J.J.B. All authors have read and agreed to the finalized version of this manuscript.

